# Hypoxia increases the methylated histones to prevent histone clipping and redistribution of heterochromatin during Raf-induced senescence

**DOI:** 10.1101/2023.10.02.560619

**Authors:** Soojeong Chang, Ramhee Moon, Sujin Yim, Dowoon Nam, Sang-Won Lee, Seunghyuk Choi, Eunok Paek, Junho K. Hur, Youhyun Nam, Rakwoo Chang, Hyunsung Park

**Affiliations:** Department of Life Science, University of Seoul, Seoul 02504, Korea; Department of Chemistry, Korea University, Seoul 02841, Republic of Korea; Department of Computer Science, Hanyang University, Seoul 04763, Korea; Department of Genetics, College of Medicine, Hanyang University, Seoul 04763, Korea; Department of Applied Chemistry, University of Seoul, Seoul 02504, Korea

**Keywords:** Hypoxia, Oncogene-induced senescence, Histone clipping, Cathepsin L, Histone methylation

## Abstract

Hypoxia increases histone methylation by inhibiting O_2_- and α-ketoglutarate-dependent histone lysine demethylases (KDMs). This study is the first to demonstrate how the hypoxic increment of methylated histones cross-talks with other epigenetic changes, such as histone clipping, and heterochromatin redistribution (senescence-associated heterochromatin foci, SAHF) found during oncogene-induced senescence (OIS). Raf activation in primary human fibroblasts IMR90 increased cathepsin L (CTSL)-mediated clipping of histone 3 (H3), H2B and H4 at H3 A21/T22, H2B T19/K20, and H4 G11/K12, respectively. Hypoxia protected H3 from CTSL by increasing histone methylation, especially at H3K23me3 without reducing the activity of CTSL. The maintenance of methylated histones is sufficient for protecting histones from CTSL, not sufficient but necessary for inhibiting SAHFs. Expression of cleaved H3 induces senescence even under hypoxia, suggesting that hypoxia disrupts this positive feedback loop of OIS by increasing histone methylation. Thus, hypoxia protects histones and chromatin from dramatic epigenetic changes by increasing histone methylation.

**Highlights:**

✓ Raf activation in primary fibroblasts increases cathepsin L-mediated cleavage of H3, H2B, and H4.
✓ Hypoxia inhibits OIS-induced histone clipping by maintaining methylated histones.
✓ Cleaved H3 induces senescence, even under hypoxia.

## INTRODUCTION

Hypoxia is known to delay senescence (1) and differentiation (2,3) but to maintain stemness (4). The molecular mechanisms by which hypoxia regulates cell fate have been explained by the specific functions of key hypoxic target genes. Hypoxia induces several hundreds of genes involved in glycolysis, angiogenesis, cell adhesion, metastasis, and proliferation. Among these genes, approximately 50% are induced by the heterodimeric transcription factor hypoxia-inducible factor (HIF)-α/β (5). It has been shown that hypoxia can directly inhibit the activity of O_2_- and α-ketoglutarate(α-KG)-dependent dioxygenases that catalyze the oxidation-dependent demethylation of histones, DNA, and RNA. Therefore, this could be a novel mechanism through which hypoxia can regulate gene expression without activating HIF-1 (6).

Compared to DNA and RNA, histones have more diverse methylation sites, that are, lysine and arginine residues, to provide highly informative combinatorial modification patterns. The five histone isoforms, H1, H2A, H2B, H3, and H4, have several arginine and lysine residues in their tails. The number of identified methylations on these residues is increasing, although methyltransferases and demethylases acting on each residue have not been identified. Theoretically, methylation levels of individual residues depend on the balance between the activities of the corresponding methyltransferases and demethylases. Because most histone demethylases are Jumonji domain-containing histone demethylases (JHDMs), which use O_2_ and α-KG as co-substrates, variations in O_2_ and α-KG concentrations in the microenvironment contribute to heterogeneity in the methylation levels of individual lysine and arginine residues (7,8).

The ENCODE project has built a platform for systematic mapping of histone modifications, chromatin accessibility, chromatin topology, and transcription along the genomic sequences of numerous tissues and cell types to provide multi-layered sequencing information (9). Trimethylation at the 4^th^ lysine of histone 3 (H3K4me3), H3K9me3, and H3K27me3 has been extensively investigated to identify their specific methyltransferases, demethylases, binding proteins, and DNA-binding sequences. H3K4me3 and H3KAc are specifically found at the transcription start sites (TSS) of actively transcribed genes and accessible euchromatin, whereas H3K9me3 and H3K27me3 are found in the promoters of repressive genes and inaccessible compact heterochromatin (10). Although causality is unclear, many histone modifications are highly related to chromatin structure and activity. The finding that each histone modification is determined by specific pairs of modifying and erasing enzymes suggests that histone modifications can be reversible depending on the proximities and activities of the enzymes (11).

Recently, the N-terminus of H3 has been found cleaved in several species and during cellular processes such as yeast sporulation, in mouse intestinal villi, in the human monocytes, in the *in vitro* neuronal differentiation of stem cells (12–14). In contrast to reversible modifications, histone clipping is irreversible if it is not actively replaced by newly synthesized histones (15–17). Histone clipping is not commonly found but is especially observed during irreversible cellular processes such as terminal differentiation and oncogene-induced senescence, suggesting that histone proteolysis plays a crucial role in determining cell fate and development. The forced expression of oncogenes such as Ras, Raf, and Myc in cultured primary cells induces rapid cellular senescence by activating the p53 and Rb pathways (18). In human melanocytic nevi, senescent cells that contain a high proportion of B-RAF V600E showed *in vivo* evidence that oncogene-induced senescence (OIS) acts as a surveillance system to protect against tumor development. Raf-induced premature senescence blocks the spread of oncogenic lesions during the early stages of tumor development (19,20). However, the OIS process also induces the expression of secreted inflammatory factors that constitute precancerous microenvironments, thus OIS has both anti- and pro-cancerous effects (21).

The OIS process increases active cathepsin L, a lysosomal cysteine protease, in the nucleus, leading to the proteolytic clipping of H3 (22,23). In addition to histone clipping, OIS is accompanied by dramatic epigenetic changes in histone modifications and chromatin topology. Several studies have shown that hypoxia attenuates senescence by regulating the DNA damage response, cell cycle, and AMPK-mTOR pathway (24–27). This study first investigated whether the hypoxia-induced increase in methylated histones affects other epigenetic changes such as histone clipping and redistribution of heterochromatin during OIS, leading to the prevention of OIS.

This study reports that during Raf-induced senescence, CTSL cleaved not only H3, but also H2B and H4. Hypoxia prevented Raf-activation from clipping H3 not by inhibiting CTSL but by increasing H3K18me3 and H3K23me3 near the cleaved site of H3. Ectopic expression of cleaved H3 partly induced senescence, even under hypoxic conditions. This study suggests that hypoxia protects histones from proteolytic cleavage by maintaining methylated histones to regulate cell fate.

## MATERIALS AND METHODS

### Cell culture

Human lung fibroblast IMR90 cells (CCL-186, ATCC, Manassas, VA, USA) were cultured in Eagle’s minimum essential medium (10-009, Corning, Glendale, AZ, USA) supplemented with 10% fetal bovine serum (FBS), 100 IU/ml penicillin, and 100 μg/ml streptomycin. Cells were cultured in humidified air containing 5% CO_2_ at 37°C. Cells were incubated under hypoxic conditions in a Forma anaerobic incubator (Model 1029, Thermo Fisher Scientific, Waltham, MA, USA) in an atmosphere of 5% CO_2_, 95% N_2_, and 0.5% O_2_.

### Generation of stable cell lines using retrovirus or lentivirus infection

IMR90 cells were infected with a retrovirus encoding ΔB-RAF:ER obtained from Dr. Martin McMahon (University of Utah, Salt Lake City, UT, USA.) to generate IMR90-ΔB-RAF:ER cells. The ΔB-RAF:ER fusion protein consists of a protein kinase domain of B-RAF (449–804 aa) and a mutant form of the hormone-binding domain of the estrogen receptor (281– 599 aa, with a single amino acid change from glycine to arginine at 525) that has been engineered to be nonresponsive to β-estradiol but retains responsiveness to 4-hydroxy-tamoxifen (4-OHT) (28). IMR90-ER, IMR90-H3.1:ER, IMR90-H3.1cs1:ER, IMR90-H3.3:ER, and IMR90-H3.3cs1:ER cells were generated by infecting IMR90 cells with retroviruses encoding ER, ER tagged H3.1, H3.1cs1, H3.3, and H3.3cs1, respectively, using the pBABE retroviral vector system and HEK293-based packaging cells (AmphoPackTM 293 cell line) (3). IMR90-ΔB-RAF:ER-M56CTSL and IMR90-ΔB-RAF:ER EV cells were generated by infecting IMR90-ΔB-RAF:ER cells with lentiviruses expressing Myc-His-tagged M56 CTSL or empty vector, respectively, using the pLenti CMV vector system. IMR90-ΔB-RAF:ER-shCTSL and IMR90-ΔB-RAF:ER-shScramble cells were generated by infecting IMR90-ΔB-RAF:ER cells with lentiviruses encoding shRNA against human CTSL (5’-GGTGGTTGGCTACGGATTTGA-3’) and scramble (control) (5’-CCTAAGGTTAAGTCGC CCTCG-3’), respectively, using the pLKO.1 lentiviral vector system (Addgene, Cambridge, MA, USA). A list of the stable cell lines generated is summarized in supplementary table S1.

### Reagents and antibodies

A list of antibodies, and reagents used are summarized in supplementary table S2 and S3.

### Senescence-associated β-Galactosidase assay

Senescence-associated β-Galactosidase (SA-β-gal) activity was measured as previously described (29). The percentage of SA-β-Gal-positive cells (blue staining) was calculated as the number of SA-β-Gal-positive cells divided by the total number of cells. Counts were performed in at least three randomly selected microscopic fields from two independent experiments to determine the mean percentage of positively stained cells.

### Quantitative Reverse Transcription-PCR (qRT-PCR)

Steady-state mRNA expression was measured using quantitative real-time PCR, as described previously (Park et al. 2013). The pairs of forward and reverse primer sequences for qRT-PCR were as follows: human cathepsin L (F) 5’ -AAACTGGGAGGCTTATCTCACT - 3’ / (R) 5’- GCATAATCCA TTAGGCCACCAT - 3’ and 18S rRNA (F) 5’ - ACCGCAGCTAGGAATAATGGAATA – 3’/ (R) 5’ - CTTTCGCTCTGGTCCGTCTT - 3’

### Immunofluorescence microscopy and Quantification of SAHF

The cells were labeled with the indicated primary antibodies and secondary antibodies conjugated to Alexa Fluor ^®^ 488 or Alexa Fluor ^®^ 546 (Invitrogen, Waltham, MA, USA) as described previously (30). The cells were mounted onto coverslips with Hoechst 33258 and observed under a Zeiss LSM800 inverted confocal microscope (Carl Zeiss, Thornwood, NY, USA), according to the manufacturer’s instructions. The primary antibodies used for immunofluorescence are summarized in supplementary table S2. The degree of SAHFs was quantified using the coefficient of variation (CV) of Hoechst or H3K9me3 intensity as described (31). Using ImageJ software that identifies Hoechst and H3K9me3 stained nuclei, the CV was calculated as the ratio of the standard deviation to the mean intensity of Hoechst or H3K9me3. The CV was presented as boxplots. The midline of the boxplot represents the median, the bottom of the box represents the 25th percentile, the top of the box represents the 75th percentile, and whiskers represent the minimum and maximum values.

### Top-down proteomic analyses

The histone pellets isolated as described above were air-dried for 10 min and resuspended in 100 μL of ddH_2_O instead of 4 M urea. The concentration of histone proteins was measured using BCA assay (23225, Thermo Fisher Scientific, Waltham, MA, USA). Proteins were aliquoted into several tubes for top-down analysis and western blot analysis, and then freeze-dried. Top-down analysis was performed using a Q Exactive HF-X mass spectrometer (Thermo Fisher Scientific) coupled to a dual online LC system based on a nanoACQUITY (Waters, Milford, MA, USA) with a dual online LC system, as previously described (32). Briefly, two analytical columns (75 µm × 100 cm, packed) and two solid-phase extraction columns (150 µm × 3 cm) packed with MAbPac particles (C8 resin, 4 µm diameter, 300-Å pore size; Thermo Fisher Scientific) were used. Each histone sample (400 ng) from four different conditions was separated using 120-min gradient (25–35% solvent B over 101 min, 35%–80% over 5 min, 80% for 10 min, and holding at 1% for 4 min). Solvent A and B comprised 0.1% formic acid in water and 0.1% formic acid in ACN, respectively. The column flow rate was 600 nl/min. The Q Exactive HF-X was tuned by setting the trapping gas pre-setting value to 0.2 in order to maintain the ultra-high vacuum range under 5.0 × 10^-11^. The eluted proteins were ionized at an electric potential of 2.7 kV and the temperature of desolvation capillary at 300℃. Full MS scans were acquired with an m/z range of 500–2000 That a resolution of 120000. The maximum ion injection time and target values for automatic gain control (AGC) were 250 ms and 5.0 × 10^6^, respectively. For multiplexed MS/MS scans, ions of the same protein but different charges (up to six charges) were simultaneously isolated and fragmented at a resolution of 60,000. Multiplexed MS/MS of an intact protein showed increase in sequence-specific fragments, and thereby increased sequence coverage. The AGC target value and maximum ion injection time for multiplexed ions were set to 5.0 × 10^6^ and 125 ms, respectively. One full MS scan and 20 targeted MS/MS scans were repeated during a 120 min run time. The number of micro-scan and normalized collision energy (NCE) were 3 and 25, respectively, and the isolation window was 0.6 Th. The LC-MS/MS data were analyzed using FLASHDeconv2D to obtain monoisotopic masses from the MS spectra and MODplus for annotating multiplexed MS/MS spectra (33,34). VisioProt-MS was used to construct a 2D feature map based on monoisotopic masses obtained from FLASHDecov2D (35).

### Competitive enzyme kinetic assay

Recombinant human Cathepsin L (rhCTSL) (Catalog # 952-CY) and Fluorogenic Peptide Substrate VII: Z-Leu-Arg-4-methyl-7-coumarylamide (Z-LR-AMC)(Catalog # ES008) were purchased from R&D Systems (Minneapolis, MN, USA). The enzyme reactions were initiated upon addition of ice-cold of hrCTSL (96pM) to the nine different concentrations of Z-LR-AMC (0, 0.5, 1, 2, 4, 8, 12, 16, 20 μM) in 100 μl of assay buffer (50 mM MES (pH 5.5), 5 mM DTT, 1 mM EDTA, and 0.005% Brij-35). Each reaction was incubated in triplicates in 96 well plates at 37℃ for 20 min to 1 hr. Using a SpectraMax iD3 Multi-Mode Microplate Reader (Molecular Devices, San Jose, CA, USA), the fluorescence changes of Z-LR-AMC were monitored in every 1 min for 20 ∼ 60 minutes at excitation and emission wavelengths of 360 nm and 460 nm (bottom read), respectively. The initial velocities of enzyme reaction (RFU/min) of three replicates (n=3) were continuously monitored assuming fast equilibrium. The velocities remained constant and linear for initial 5 min. The initial velocity at each starting concentration of substrate was determined using slope of linear regression. To determine the Km value of recombinant human CTSL for a substrate, Z-LR-AMC, initial velocity was fitted to the Michaelis–Menten model in GraphPad Prism 8.0 (GraphPad Software Inc., San Diego, CA, USA). For the competitive inhibition assay, naïve or methylated H3 peptides (100 nM or 400 nM) were added to Z-LR-AMC (1, 4, 12, 20 μM) together with 96 pM of rCTSL. The initial velocities were determined as described above. Ki values were determined according to the competitive inhibition model, KmObs (observed Km) = Km×(1+[I]/Ki), in GraphPad Prism 8.0 (GraphPad Software Inc., San Diego, CA, USA) with a nonlinear regression function. All kinetic assays were performed in triplicates.

### Molecular dynamics simulations

The molecular dynamics (MD) simulations were performed with OpenMM software and CHARMM36m force field (36,37). The crystal structure of the C25A mutant of Cathepsin L (CTSL) in complex with Gln-Leu-Ala peptide (PDB id 3K24) was used to model the CTSL systems. The mutated residue Ala25 was modified back to Cys25 and only one monomer of the original dimer structure was used in the simulation because the active site was far from the interfacial region between the monomers. The protonation states of 71, 109, 9, 50, 119, 140, 163 residues in the enzyme were calculated with Propka3.1 in the experimental condition of pH 5.5. In the MD simulations, the two different 6-residue fragments (H3_19-24_) of H3 were used as the substrates: naïve H3_19-24_ and trimethylated H3_19-24_K23me3 (38,39). The position of first three residues in the substrate was taken from the X-ray structure and the unknown location of the last three residues (Thr-Lys-Ala) in the substrates was initially predicted by AutoDock Vina (40,41), which was later confirmed as the most stable conformation by scanning the energy landscape along the dihedral angles in the backbone of the last three residues. To prevent additional interactions with the cleaved termini, the methyl group was attached to both the N- and C-terminal of both enzyme and tail fragments. The simulation box size was initially set to 80 x 80 x 80 Å^3^. Initial system settings such as physiological conditions (KCl 0.15 M), terminal and hydrogen patches, water solvation, and periodic boundary conditions were performed with CAHRMM-GUI (42–44). Each system was first energy minimized for 5000 steps and then was equilibrated for 500 ps in NVT and NPT ensembles with gradual heating up to 310 K at 1 bar. Then, the equilibrium 100 ns MD simulation for each system was then performed in the NPT ensemble (1 bar and 310K) without any restraints. Both the Lennard-Jones and the real part of the electrostatic interactions used the cutoff radius of 10 Å and the reciprocal part of the electrostatic interaction was treated by using the particle mesh Ewald (PME) summation method (45,46). The thermostat and barostat used in the simulation were Langevin dynamics and Monte Carlo methods, respectively. The first 15 ns was not equilibrated, and the rest 85 ns trajectory was used for the results presented in this study.

### In vitro CTSL reaction

*In vitro* proteolytic reaction of human recombinant cathepsin L (50 pg) was performed with recombinant histones (200 ng) at 37℃ for 30 min in 20 μl of reaction buffer composed of 50 mM MES pH 5.5 containing 5 mM DTT, 1 mM EDTA, and 0.005% Brij-35. To obtain endogenous histones, we isolated the nuclear fraction from IMR90 cells cultured under hypoxia or normoxia, as previously described (47). For each *in vitro* proteolytic reaction, nuclear fraction from 30,000 nuclei were treated with 1 ng or 2 ng of recombinant human cathepsin L in 20 μl of reaction buffer for 30 min at 37℃.

### Analysis of cellular cathepsin L activity

To measure the activity of CTSL in live cells, we used the Magic Red Cathepsin L Detection Kit (Immunochemistry Technologies, Bloomington, MN, USA). Briefly, 5,000 suspended cells were loaded with Magic Red Cathepsin L reagent containing a CTSL substrate, N-carbobenzyloxy-Phe-Arg-MR-Arg-Phe (MR-FR2), for each enzyme reaction. The fluorescence was then measured (Ex 590 nm, Em 630 nm) using a kinetic mode of SpectraMax® iD3 (Molecular Devices) at 37℃ for 1 h using the bottom-read function. The enzyme reaction velocity (RFU/min) was determined using GraphPad Prism software (GraphPad Software, Version 8.0) using a linear regression function.

### Assay for transposase-accessible chromatin (ATAC)

We constructed an assay for transposase-accessible chromatin (ATAC) using a Tn5-Nextera DNA library prep kit (Illumina, San Diego, CA, USA), following the manufacturer’s protocol (48). Briefly, 50,000 nuclei from each group of cells were incubated in 50 µL of transposition mix containing transposase for 30 min at 37℃. Transposed DNA was purified using the MinElute PCR Purification Kit (QIAGEN, Germantown, MD, USA). DNA fragments were then amplified by PCR using Nextera index primer 1 and 2 (Illumina) and produced by PCR amplification (10–13 cycles) of tagmented DNA using a NEB Next High-Fidelity 2× PCR Master Mix (New England Biolabs, Ipswich, MA, USA). DNA fragments were then purified using the MinElute PCR Purification Kit and eluted in 10 µL elution buffer. Libraries were assessed for fragment length distribution on a Bioanalyzer High-Sensitivity DNA Chip (Agilent Genomics, Santa Clara, CA, USA).

### Assay of transposase-accessible chromatin with visualization (ATAC-see)

Cells were fixed with 1% formaldehyde for 10 min at room temperature, permeabilized with cell lysis buffer (10 mM Tris–Cl, pH 7.4, 10 mM NaCl, 3 mM MgCl2, 0.01% Igepal CA-630) for 10 min, and then washed with PBS. The fluorescent Atto-590 attached Tn5 (Atto-590 Tn5) was added to cells on the slide and incubated for 30 min at 37℃. After the transposase reaction, the slides were washed with PBS containing 0.01% SDS and 50 mM EDTA for 15 min, three times at 55℃. The cells were then co-stained with either H3K9me3 or H3K4me3 antibodies and subsequently mounted with Hoechst 33258 (49). The fluorescence intensity of the Atto-590 fluorescence signal in the nucleus was measured using ImageJ software and is presented as boxplots. The midline of the boxplot represents the median, the bottom and top of the box represent the 25th and 75th percentiles, respectively, and the whiskers represent the minimum and maximum values.

## RESULTS

### Hypoxia-induced protection of histone clipping during OIS

We generated IMR90-ΔB-RAF:ER cells, primary human lung fibroblast IMR90 cells that express a constitutively active form of ΔB-Raf mutant fused with the ligand-binding domain of the human estrogen receptor (ER) (28,50). ΔB-RAF:ER fusion protein was stabilized only upon treatment with the ER antagonist 4-OHT (Fig. 1A). We exposed IMR90-ΔB-RAF:ER cells to hypoxic conditions (0.5% O_2_) during 4-OHT treatment and confirmed that 4-OHT also induced ΔB-RAF:ER fusion protein under these conditions and that hypoxia increased the levels of HIF-1α, a hypoxia master transcription factor (Fig. 1A and 1B). Raf activation also increased the activity of SA-β gal and senescence-associated heterochromatin foci (SAHF), hallmarks of OIS (Fig. 1C and 1D). SAHF was visualized by immunostaining for H3K9me3 (trimethylated 9^th^ lysine of histone 3) and heterochromatin protein (HP)-1β which constitute heterochromatin (Fig. 1D and 1E). Immunostaining for H3K27me3, which localizes to the boundary of SAHF, also confirmed that Raf activation induced SAHF (Fig. 1F). Interestingly, hypoxic condition prevented active Raf from increasing SA-β-gal activity and SAHF.

**Figure 1.**
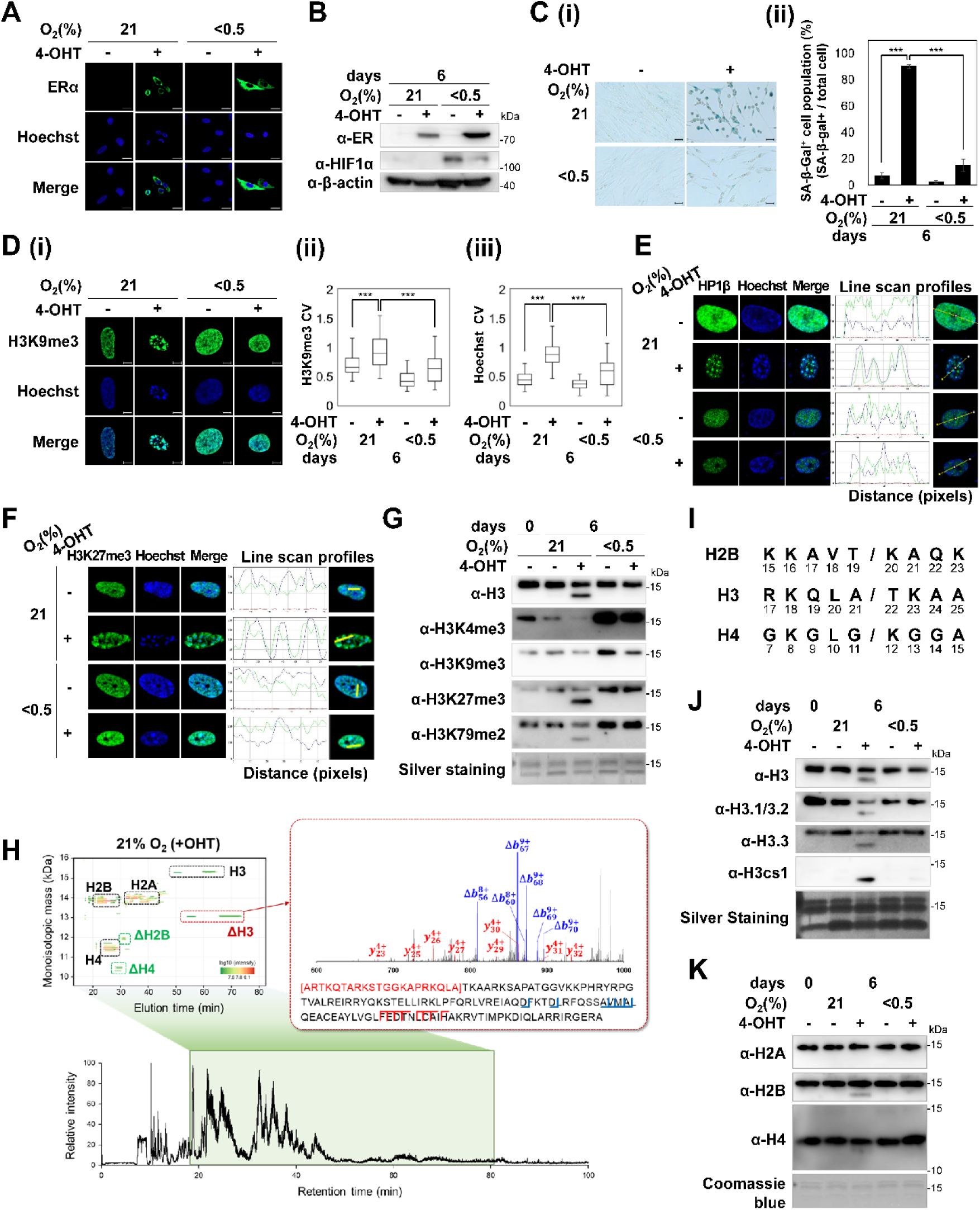
Hypoxic inhibition of histone clipping in 4-OHT treated IMR90-ΔB-RAF:ER cells. IMR90-ΔB-RAF:ER cells were treated (+) with 4-OHT (100 nM) for six days under normoxic (21% O_2_) or hypoxic (0.5% O_2_) conditions. (A) Confocal microscopic images obtained following ERα antibody and Hoechst staining. (B) Western blot analysis with the indicated antibodies. β-actin was used as a loading control. (C) (i) Representative images of the SA-β-gal assays. (ii) Mean and S.D. of SA-β-gal-positive cells from six randomly selected microscopic fields from three independent experiments. SA-β-gal-positive cells among 180– 890 cells per field were counted. (D) (i) Confocal microscopic images of nuclei stained with the H3K9me3 antibody and Hoechst. (ii-iii) SAHFs were measured using the coefficient of variation (CV) of nuclear images stained with H3K9me3 or Hoechst (details in Methods). The values on the y-axis represent the CV of 55–60 nuclear images from two independent experiments. (E-F) Confocal microscopic images of nuclei stained with the HP1β antibody (E) or H3K27me3 antibody (F) and Hoechst. The intensities of the images on the yellow straight lines were quantified using the ImageJ software. Hoechst (purple line), HP1β (green line), and H3K27me3 (green line). (G) Western blot analysis using the indicated antibodies. Silver staining was used as loading control. (H) Base peak chromatogram reconstructed from the mass spectra of LC-MS/MS data using histone sample from IMR90-ΔB-RAF:ER cells treated with 4-OHT under normoxic conditions. A 2D display (monoisotopic masses vs elution time) of intact histones measured in LC-MS/MS experiments (upper left). An annotated MS/MS spectrum labeled for cleaved N-terminal (Δb ions, blue) and C-terminal (y-ions, red) fragments shows a truncated H3 (upper right). (I) Histone cleavage sites (J-K) Western blot analysis using the indicated antibodies. Silver staining (J) and Coomassie Brilliant Blue staining (K) were used as loading controls. \*\*\**p* < 0.001 by Students’ t-test.

Upon Raf activation, cleavage of histone 3 (H3) was observed (Fig. 1G). H3 antibodies specific for H3K27me3 and H3K79me2 also detected smaller cleaved fragments of H3 in OIS cells, whereas antibodies against H3K4me3 and H3K9me3 failed to do so, suggesting that OIS resulted in the cleavage of the peptide bond between the 9^th^ lysine and 27^th^ lysine of H3. Hypoxic conditions prevented OIS from clipping H3 but also increased the methylated histones, H3K4me3, H3K9me3, H3K27me3, and H3K79me2 (6). Top-down proteomics analyses employing the multiplexed targeted MS/MS method further showed that OIS induced the cleavage of not only H3, but also H2B and H4 at H3 A21/T22, H2B T19/K20, and H4 G11/K12, respectively, however no cleavage of H2A was observed (Fig. 1H and 1I). Furthermore, top-down proteomics data showed that hypoxia prevented OIS-induced cleavage of H2B, H3, and H4 (Fig. S1A and S1B). Western blot analyses using anti-H3.1/3.2 and anti-H3.3 antibodies confirmed that OIS induced cleavage of H3.1/3.2 and H3.3 isoforms. Cleaved H3 was also detected using the H3cs1 antibody, which recognizes the H3 tail beginning at amino acid T22 (Fig. 1J). Antibodies specific for H2B or H4 also detected cleaved histones that migrated faster (Fig. 1K). We confirmed that, under hypoxic conditions, Raf activation failed to induce cleavage of H2B, H3, and H4. Neither doxorubicin, a senescence inducer, nor replicative senescence cleaved H3 (Fig. S2A to S2D). Raf activation in the cancer cell lines HT29 and HEK293 resulted neither in cleavage of H3 nor in induction of senescence (Fig. S2E and S2F). These results suggested that histone clipping is a specific phenotype of OIS.

### Hypoxic protection of histone clipping not by CTSL inhibition

We examined whether cathepsin L (CTSL) cleaves H2B, H4, and H3. Treatment with E64, a cysteine protease inhibitor specific for cathepsin B, H, and L, or knockdown of CTSL prevented OIS-induced clipping of not only H3, but also H2B and H4, indicating that CTSL cleaves H2B and H4 during OIS (Fig. 2A and 2B). We also investigated whether hypoxia hinders CTSL activation during OIS. Upon Raf activation, the mRNA and protein levels of CTSL significantly increased under both normoxia and hypoxia (Fig. 2C and 2D). OIS resulted in an increase in the amount of the processed mature form of CTSL (19 kDa) in the nuclei (Fig. 2E to 2G). Using MR-FR_2_, a fluorescence-labeled membrane-permeable synthetic substrate for CTSL (51), we found that hypoxia did not reduce the total cellular activity of CTSL (Fig. 2H). We constructed IMR90-ΔB-RAF:ER-M56CTSL cells that are IMR90-ΔB-RAF:ER cells, additionally infected with M56-CTSL-Myc-His which is a constitutively active mutant of CTSL exclusively located in the nucleus, to test whether hypoxia decreases the catalytic activity of M56-CTSL (Fig. 2E, 2I, and 2J) (52). Ectopic expression of M56-CTSL in nuclei similarly increased CTSL activity measured using MR-FR_2_ in both normoxic and hypoxic cells (Fig. 2K), however M56-CTSL significantly reduced the levels of intact H3 and H2B only under normoxia but not under hypoxia (Fig. 2L), suggesting that hypoxia hinders histone clipping during OIS, not by inhibiting CTSL activity.

**Figure 2.**
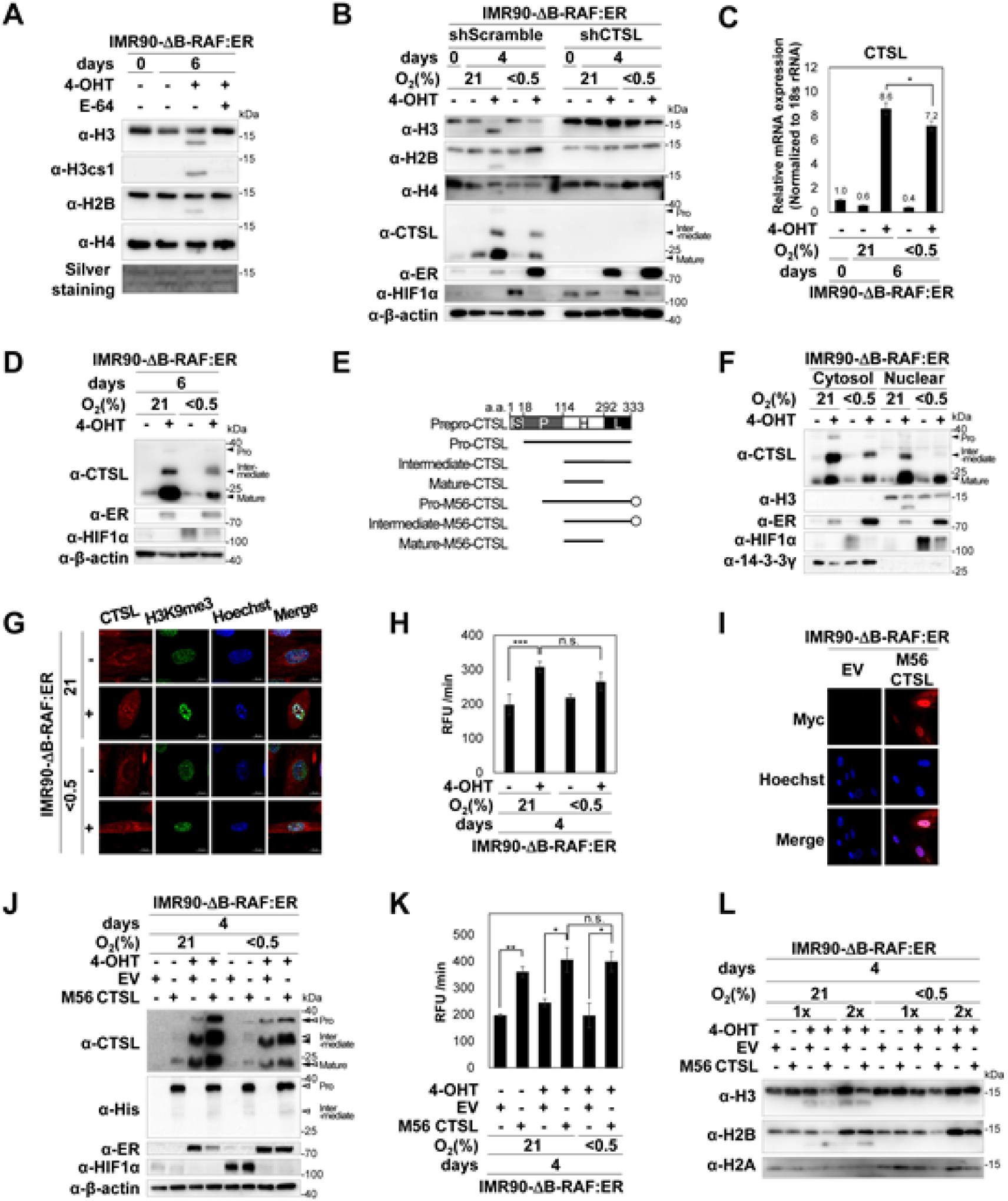
Cathepsin L-dependent histone clipping. (A-B) Western blot analysis using histone antibodies. (A) IMR90-ΔB-RAF:ER cells were treated with 4-OHT in the presence of E64 (10 μM) as indicated. (B) IMR90-ΔB-RAF:ER-shScramble cells and IMR90-ΔB-RAF:ER-shCTSL cells, which are IMR90-ΔB-RAF:ER cells additionally infected with lentivirus encoding shScramble or shCTSL, respectively. (C) Quantitative RT-PCR of CTSL mRNA of IMR90-ΔB-RAF:ER cells treated as indicated. The values on Y-axis are means and S.E. of CTSL mRNA normalized using levels of 18s rRNA from three independent experiments. (D) Western blot analysis using CTSL antibody. Pro-CTSL, intermediate-CTSL, and mature-CTSL are shown. (E) Schematic representation of processed forms of CTSL. The full-length CTSL is composed of distinct domains: a signal peptide (S), pro-domain (P), heavy chain (H), and light chain (L) (75). The deletion mutant M56-CTSL-Myc-His fusion protein translated from internal AUG located at 56^th^ methionine. Open circles indicate Myc-His-tag at C-terminus. (F) Western blots of the cytosolic and nuclear fractions of IMR90-ΔB-RAF:ER cells treated as indicated. The 14-3-3γ and H3 were used as loading controls for cytosolic and nuclear fractions, respectively. (G) Confocal microscopic images of cells stained with CTSL and H3K9me3 antibodies and Hoechst. (H) Quantification of CTSL enzyme activity in IMR90-ΔB-RAF:ER cells incubated with the CTSL substrate MR-FR2 (3.6 μM) for 1 h (details in Methods). The values indicate the means and S.D. of the reaction velocity (RFU/min) from three independent assays. (I-L) IMR90-ΔB-RAF:ER-M56CTSL cells that are IMR90-ΔB-RAF:ER cells additionally infected with M56-CTSL-Myc-His or empty vector. (I) Confocal microscopic images of cells stained with Myc antibody and Hoechst. (J) Western blot analysis using the indicated antibodies. Closed triangles indicate endogenous CTSL bands and open triangles indicate M56CTSL bands. (K) Quantification of CTSL enzyme activities in IMR90-ΔB-RAF:ER-M56CTSL cells that were incubated with the CTSL substrate MR-FR2 (3.6 μM) for 1 h (details in Methods). The values indicate the means and S.D. of the reaction velocity (RFU/min) from three independent assays. (L) Western blot analysis using the indicated antibodies. H2A was used as a loading control. 2x indicates that twice more amount of cell lysates was loaded compared to 1x. \**p* < 0.05, \*\**p* < 0.01, and \*\*\**p* < 0.001 by Students’ t-test; ns, not significant.

### Hypoxia and methylated histones

Hypoxia limited the catalytic activities of O_2_-dependent Jumonji domain-containing histone lysine demethylases (JHDMs)/KDMs (6). Hypoxia and JIB-04, a pan-inhibitor of JHDMs/KDMs, increased the levels of methylated histones, such as H3K4me3, H3K9me3, and H3K27me3 (Fig. 3A). Similar to hypoxia, JIB-04 and other pan-inhibitors of JHDMs/KDMs, such as dimethyloxalylglycine (DMOG), ciclopirox (CPX), and deferiprone (DFP) also prevented OIS-induced cleavage H3, H2B, and H4, suggesting that methylated histones could lose their substrate susceptibility to CTSL (Fig. 3B and 3C). Neither JIB-04 nor DMOG reduced CTSL levels (Fig. 3D). *In vitro* proteolytic reaction using recombinant histones and recombinant CTSL revealed that E64 inhibited CTSL-mediated to cleavage of H3 and H2B, whereas JIB04 failed to directly prevent CTSL-mediated cleavage of unmodified recombinant histones, because JIB04 cannot increase the methylation of naïve recombinant histones *in vitro* (Fig. 3E and 3F). Instead of using recombinant histones, we used nuclear fractions isolated from IMR90 cells cultured under either normoxia or hypoxia for *in vitro* CTSL reactions. We found that histones in nuclear fractions isolated from hypoxic cells were more resistant to CTSL than those isolated from normoxic cells, because histones isolated from hypoxic cells were more methylated than those isolated from normoxic cells (Fig. 3A and 3G). These results suggest that methylation makes histones resistant to CTSL and that hypoxia, JIB-04, and other inhibitors of histone demethylases protect histones from CTSL by increasing their methylation.

**Figure 3.**
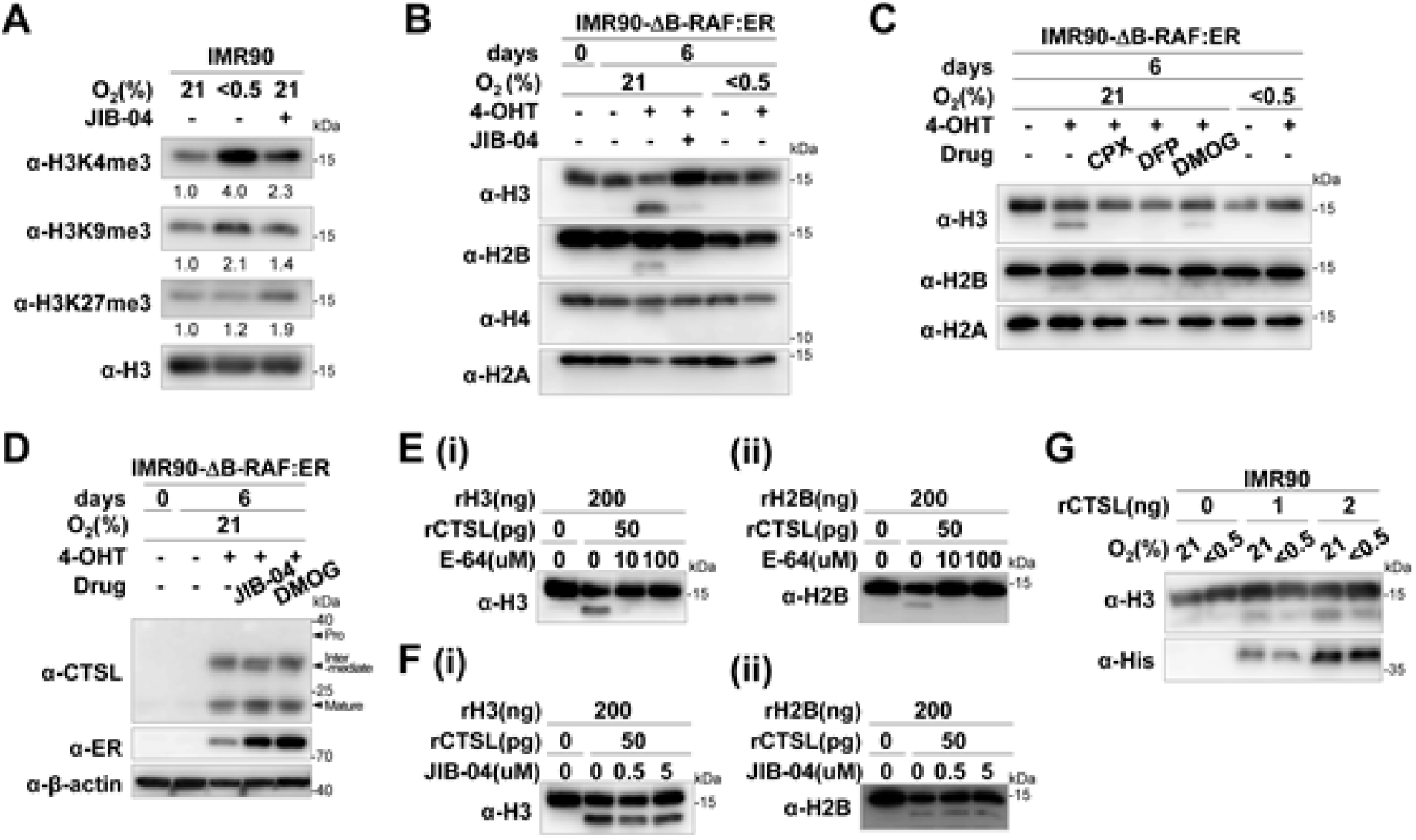
*In vitro* proteolytic reaction using recombinant CTSL and histones. (A) Western analysis of IMR90 cells treated with JIB-04 (0.5 μM) and hypoxia for 6 days. The numbers represent the relative band intensities. H3 was detected as a loading control. (B) Western blot analysis using the indicated antibodies. (C) Western blot analysis of histone extracts of IMR90-ΔB-RAF:ER cells treated with CPX (10 μM), DFP (250 μM), and DMOG (0.5 mM) as indicated. (D) Western analysis of IMR90-ΔB-RAF:ER cells treated with JIB-04 (0.5 μM) and DMOG (0.5 mM) as indicated. (E-F) Western blot analysis of *in vitro* proteolytic reactions using the recombinant histone H3 and H2B together with human recombinant CTSL (rCTSL) in 20 μl of reaction buffer in the presence of E64 (10, 100 μM) (E) or JIB-04 (0.5, 5 μM) (F). Intact and cleaved fragments of histones were visualized by western blot analysis using H3 and H2B antibodies. (G) Western blot analysis of *in vitro* proteolytic reaction using the indicated amounts of rCTSL (0, 1, 2 ng) and nuclear fractions from 30,000 nuclei of IMR90 cells cultured under normoxia or hypoxia for 6 days in 20 μl of reaction buffer.

### Inhibition of CTSL-mediated proteolysis by methylation of H3K23 and H3K18

H3 has lysine residues at the 18^th^ and 23^rd^ positions close to the CTSL clipping site (H3A21/T22) (Fig. 4A). Although H3K23me1/2/3 and H3K18me1/2/3 have been identified in Tetrahymena and human cancer cells, their methyltransferases and demethylases have not been identified (53–55). We found that both hypoxia and JIB-04 increased H3K23me3, H3K23me2, H3K18me3, and H3K18me2 (Fig. 4A), and that OIS resulted in the reduction of all of them under normoxia, but not under hypoxia or in the presence of JIB-04 (Fig. 4B). We found that acetylation at H3K18 and H3K23 residues was also diminished in OIS cells, but not under hypoxia, and that unlike methylation, JIB-04 failed to maintain acetylation during OIS (Fig. 4C). These results suggest that (i) during OIS, methyl and acetyl groups were removed from histones; (ii) hypoxia protects methylated histones as well as acetylated histones; and (iii) among these modifications, methylation of H3K18 and H3K23 prevents CTSL from clipping histone H3.

**Figure 4.**
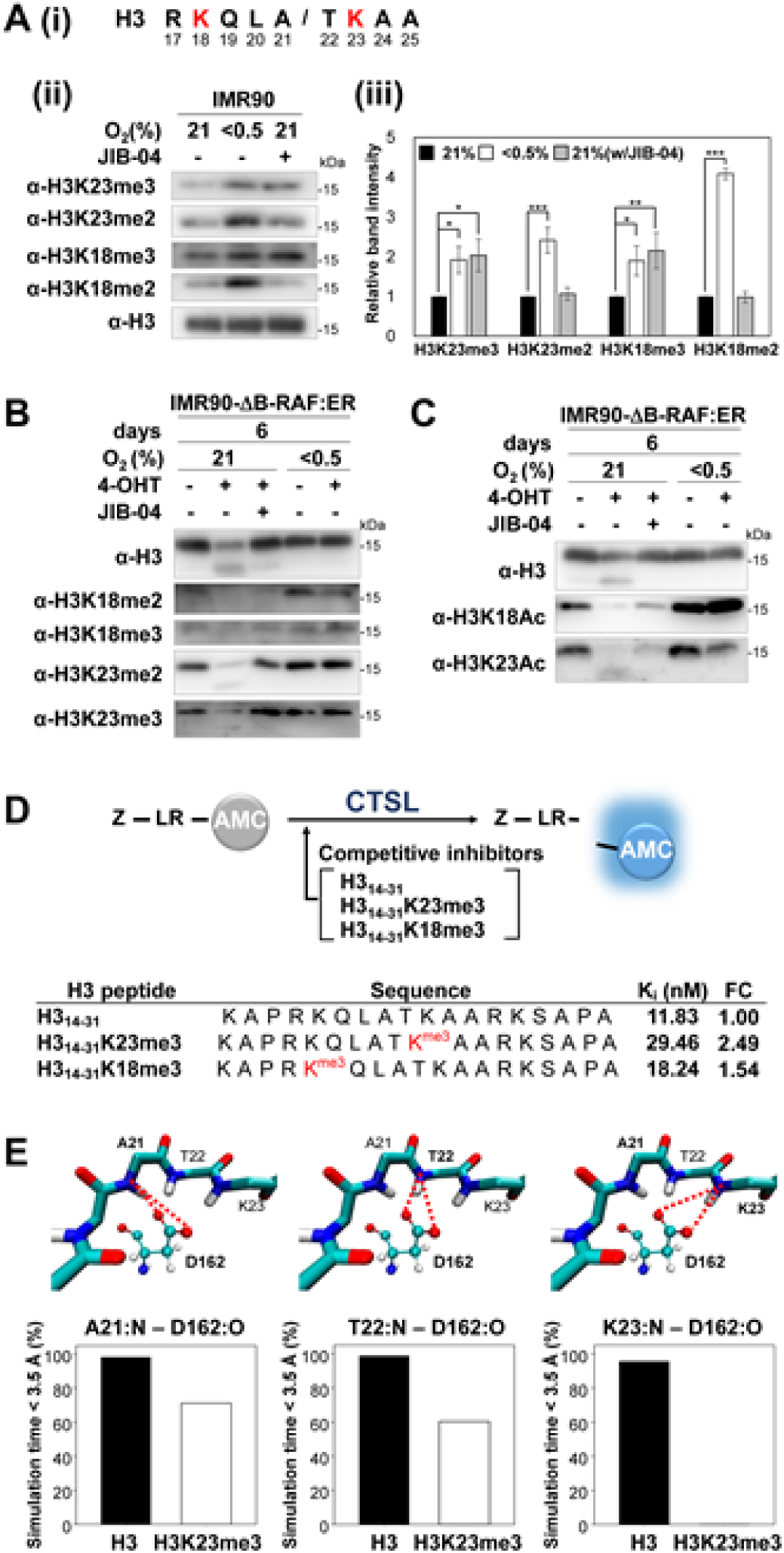
CTSL proteolysis inhibition by methylation of H3K23 and H3K18. (A) (i) Lysine residues and CTSL cleavage sites of H3. (ii) Western blot analysis of IMR90 cells exposed to hypoxia (0.5% O_2_) or JIB-04 (0.5 μM) for 4 days. (iii) The y-axis indicates the mean and S.D. of band intensities of the indicated methylated H3 normalized by H3 intensity. n=3 to 7, \**p* < 0.05, \*\**p* < 0.01, and \*\*\**p* < 0.001 by Students’ t-test; ns, not significant. (B-C) Western blot analysis of IMR90-ΔB-RAF:ER cells treated as shown. (D) Schematic diagram of the *in vitro* rCTSL activity analyses with the histone peptides as competitive inhibitors. Ki values of naïve or methylated H3 peptides for substrate Z-LR-AMC by rCTSL. The Ki values were determined using initial velocity of three replicates and nonlinear regression fit with GraphPad Prism software from a representative single experiment among two independent experiments. (E) Molecular dynamics simulations. (i) Representative simulation snapshots of D162 of CTSL and H319-24. C, O, H, and N atoms are shown in cyan, red, white, and blue colors, respectively. The red lines represent the hydrogen bonds between the hydrogen donors (N in A21, T22 and K23 of H319-24) and the hydrogen acceptors (O in D162 of CTSL): (left) A21:N – D162:O, (middle) T22:N – D162:O, and (right) K23:N – D162:O. (ii) The percentage of the hydrogen bonding between the three hydrogen-bonding pairs for the H319-24 and H319-24K23me3 (H3K23me3) systems. The percentage of the hydrogen bonding was calculated by counting the number of simulation snapshots, in which the distance between the corresponding hydrogen donor and acceptor was within 3.5 Å, from the equilibrated simulation trajectory (85 ns).

We investigated whether methylation at K18 and K23 endows resistance of H3 to CTSL. We performed enzyme kinetic analyses using recombinant human CTSL and its substrate, Z-LR-4-methyl-7-coumarylamide (Z-LR-AMC) (Fig. 4D) (56). As competitive inhibitors for CTSL, we used naïve histone3 peptide (H3_14-31_), which includes the 18 amino acids from the 14^th^ lysine to the 31^st^ alanine of H3, and other modified H3 peptides, that is, K23 tri-methylated (H3_14-31_K23me3), K18 tri-methylated (H3_14-31_K18me3) histone3 peptides as shown in Fig. 4D. The estimated Ki values indicate that H3_14-31_K23me3 has the highest Ki value (nM), suggesting that it is the worst substrate for CTSL among competitive inhibitors and that H3_14-31_K18me3 is a worse substrate than the H3_14-31_ peptide (Fig 4D, Fig S3A and S3B). We have also investigated the hydrogen bonding between CTSL and H3_19-24_ to understand the molecular origin of the different substrate susceptibility of naïve H3 and H3K23me3 substrates. Molecular dynamics simulations first revealed that D162 residue in catalytic domain of human CTSL had a crucial role in stabilizing the naïve H3 backbone (Fig. 4E). In the case of H3_19-24_, at least one of the oxygens of the deprotonated carboxyl group in D162 was interacting with the backbone of A21T22K23 by hydrogen bonds. On the other hand, the proportion of hydrogen-bonding of D162 with the H3_19-24_K23me3 was significantly reduced compared to that with the H3_19-24_. These results that tri-methylation at H3K23 confers resistance to CTSL and that hypoxia increases tri-methylation of H3K23 suggest that hypoxia can reduce substrate susceptibility of the H3 to CTSL by maintaining high H3K23me3 levels.

### H3K23me3, H3K18me3, and senescence

Immunostaining revealed that H3K23me3 and H3K18me3 were co-localized in the DNA condensed regions of SAHF. H3K23me2 was located at the boundary of SAHF, similar to H3K27me3 (55), whereas H3K23Ac did not overlap with SAHF in OIS cells (Fig. 5A, 5B, S4A and S4B). Consistent with previous findings, these observations indicate that H3K23 and H3K18 methylations co-localize with heterochromatin (57). Hypoxic cells were treated with adenosine dialdehyde (AdOx), an inhibitor of histone methyltransferase, to reduce H3 methylation (Fig. 5C and S4C). We found that AdOx treatment reduced H3K18me3 and H3K23me3 even under hypoxia, and that in the presence of AdOx, Raf activation resulted in the cleavage of H3 even under hypoxia. Furthermore, treatment with both AdOx and DZNep, another methyltransferase inhibitor, reduced hypoxic effects that prevented Raf activation from increasing SAHF (Fig. 5D) and SA-β-gal (Fig. 5E). These results suggest that increased histone methylation is necessary for hypoxia to prevent OIS from inducing senescence phenotypes, that is histone clipping, SAHFs, and SA-β-gal activation.

**Figure 5.**
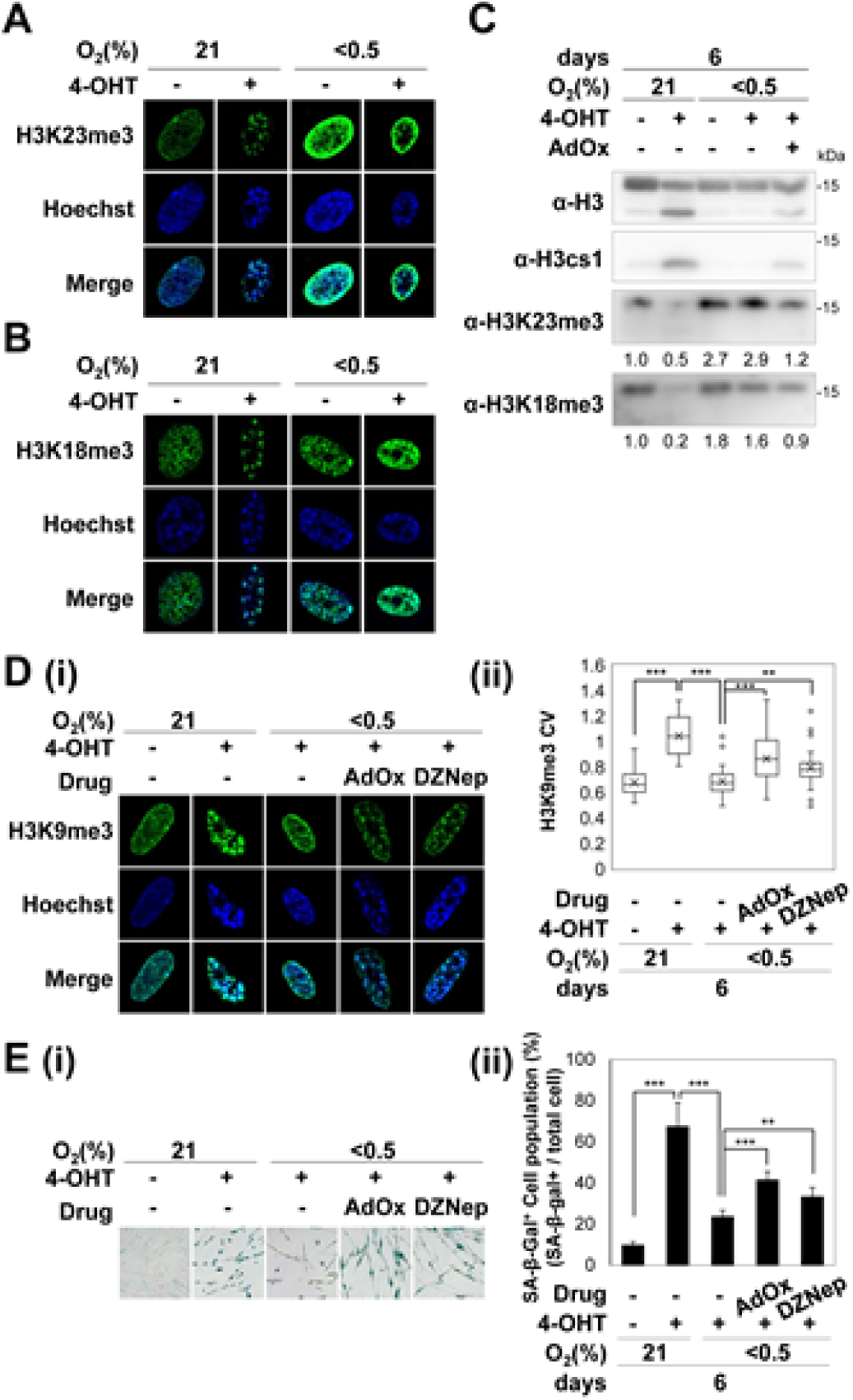
Inhibition of methyltransferase and OIS under hypoxia. IMR90-ΔB-RAF:ER cells were treated for six days. (A-B) Confocal microscopic images of nuclei stained with antibodies against H3K23me3 (A), H3K18me3 (B), and Hoechst. (C-E) IMR90-ΔB-RAF:ER cells were treated with the pan-inhibitors of methyltransferase AdOx (50 μM) and DZNep (10 μM) for 6 days. (C) Western blot analysis. The numbers represent the relative band intensities of methylated H3 normalized to the total H3 band intensities. (D) (i) Confocal microscopic images of nuclei stained with the H3K9me3 antibody and Hoechst (ii) CV boxplots of images stained with H3K9me3. H3K9me3 intensities of total 30–40 nuclear images were counted. (E) (i) Representative images of the SA-β-gal assays. (ii) Mean and S.D. of SA-β-gal-positive cells from three to five randomly selected microscopic fields. SA-β-gal-positive cells among 180–600 cells per field were counted. \**p* < 0.05, \*\**p* < 0.01, and \*\*\**p* < 0.001 by Students’ t-test; ns, not significant.

### Histone clipping. SAHFs and chromatin accessibility

To examine whether histone clipping increases chromatin accessibility, we performed an assay of transposase-accessible chromatin with visualization (ATAC-see) using a hyperactive Tn5 transposase-conjugated fluorophore adaptor (Atto590-Tn5). ATAC-see signals co-localized with the euchromatin marker H3K4me3, but not with the repressive marker H3K9me3 (Fig. 6A) (49). OIS increased the ATAC-see signals under normoxia, however under hypoxia, ATAC-see signals were concentrated in several nuclear foci, and the overall signal was much less. These results suggest that hypoxia maintains chromatin inaccessibility, even after Raf activation. Transposed DNA was amplified by PCR, and the length distribution of the PCR fragments was analyzed on a Bioanalyzer. The size distribution of the PCR fragments had a periodicity of approximately 200 bp. Raf activation reduced the sizes of PCR fragments in both normoxic and hypoxic cells, but the transposed DNA isolated from hypoxic cells was distributed in larger sizes than that in normoxic cells (Fig. 6B and S5A). These observations suggest that the OIS process increases the accessibility of chromatin under normoxia but decreases it under hypoxic conditions. Treatment with JIB-04 and E64 did not significantly prevent Raf-activation from increasing chromatin accessibility compared to hypoxia, although both prevented histone clipping (Fig. 6B). Consistently, we found that treatment with E64 and JIB-04 failed to prevent OIS from increasing SA-β-gal and SAHF levels (Fig. 6C and 6D). These results suggest that the histone clipping alone is not sufficient for increasing chromatin accessibility, SAHF formation, and SA-β-gal during OIS. Altogether, our findings in Fig. 5D, 5E, 6C and 6D suggest that increased methylation of histones is necessary, but not sufficient for hypoxia to prevent OIS from increasing SAHF and chromatin accessibility.

**Figure 6.**
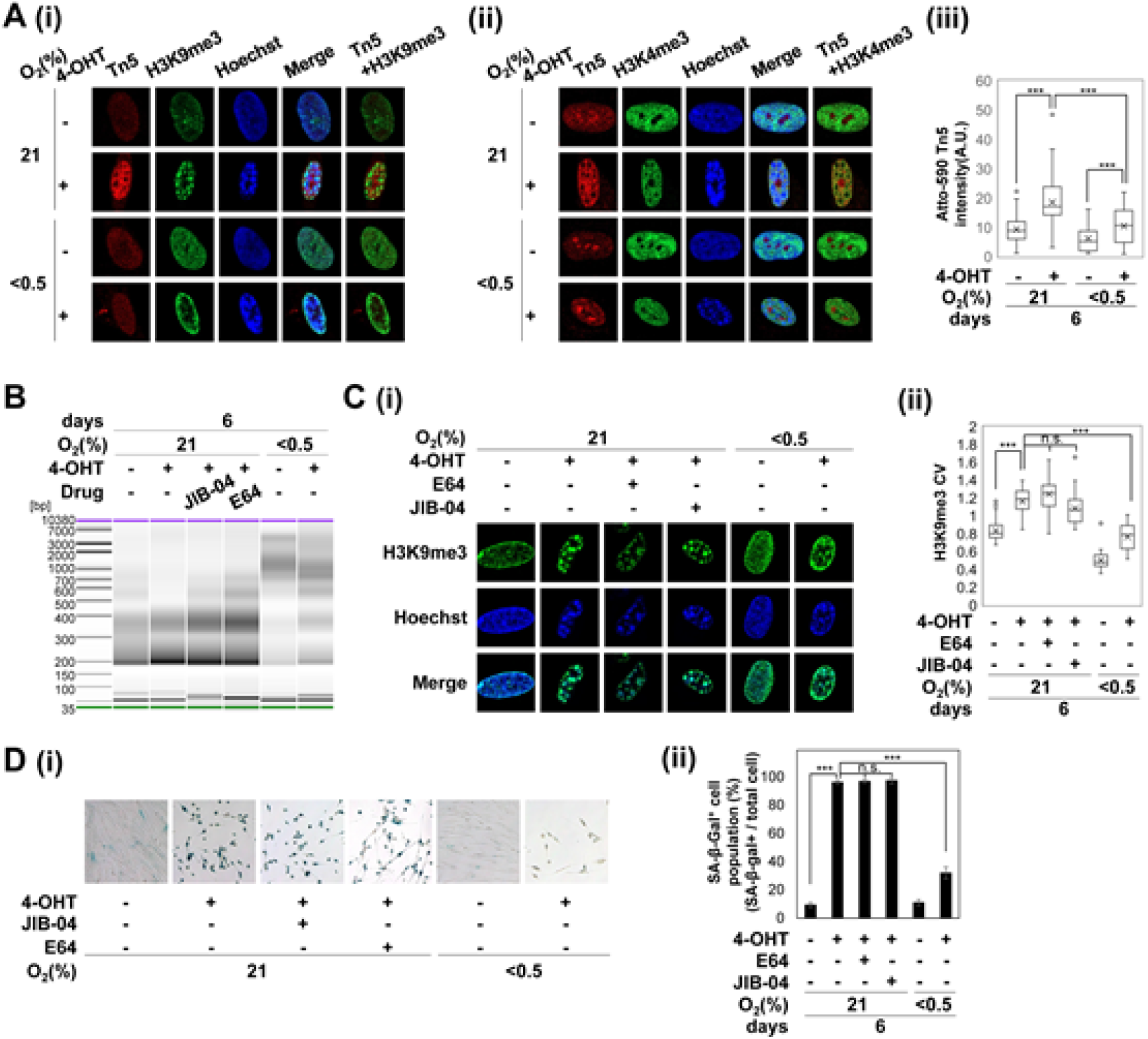
Changes in chromatin accessibility during OIS. (A) Confocal microscopic images of IMR90-ΔB-RAF:ER cells stained using Atto-590 Tn5, H3K9me3 (i), H3K4me3 antibodies (ii) and Hoechst. IMR90-ΔB-RAF:ER cells were treated as indicated for 6 days before staining. (iii) Quantification of Atto-590-Tn5 that incorporated into DNA by using ImageJ program. The intensities of Atto-590 signals in 35–50 nuclear images were counted from three independent experiments (details in Methods). (B) Fragment sizes for amplified ATAC as determined by Bioanalyzer. The data shown are from a single representative experiment out of three independent experiments shown in fig. S5A. (C) (i) Confocal microscopic images of H3K9me3 of IMR90-ΔB-RAF:ER cells treated as indicated with JIB-04 (0.5 μM) and E64 (10 μM) for 6 days. (ii) CV boxplots of images stained with H3K9me3 antibody. H3K9me3 intensities of total 28–48 nuclear images were counted. (D) (i) Representative images of SA-β-gal assay. (ii) Mean and S.D. of SA-β-gal-positive cells from three to five randomly selected microscopic fields. SA-β-gal-positive cells among 170–460 cells per field were counted. \**p* < 0.05, \*\**p* < 0.01, and \*\*\**p* < 0.001 by Students’ t-test; ns, not significant.

### Forced expression of cleaved H3 and senescence

In order to investigate the effects of H3 clipping on senescence, we ectopically expressed H3.1 and H3.3 isoforms and their cleaved forms in IMR90 primary cells. We generated IMR90-H3.3:ER and IMR90-H3.3cs1:ER cells that are IMR90 cells infected with retroviruses that encode H3.3:ER and H3.3cs1:ER, respectively. H3.3cs1:ER is a fusion protein of the N-terminus truncated H3.3 isoform (22 aa to end) and ER (Fig. 7A and S5B). In response to 4-OHT, H3.3cs1:ER and H3.3:ER were exclusively expressed in the nucleus (Fig. 7B). We found that in the absence of Raf induction, forced expression of cleaved H3 was sufficient to induce senescence, even under hypoxic conditions. Compared to H3.3:ER, H3.3cs1:ER further increased SAHF formation and SA-β-gal activity even under hypoxic conditions (Fig. 7C and 7D). Similarly, H3.1cs1:ER also increased both SAHF and SA-β-gal activity, even under hypoxia (Fig. S5C to S5E). The forced expression of either H3.3cs1 or H3.1cs1 induced senescence, even under hypoxia, suggesting that hypoxia breaks this positive feedback circle of OIS by increasing methylated histones.

**Figure 7.**
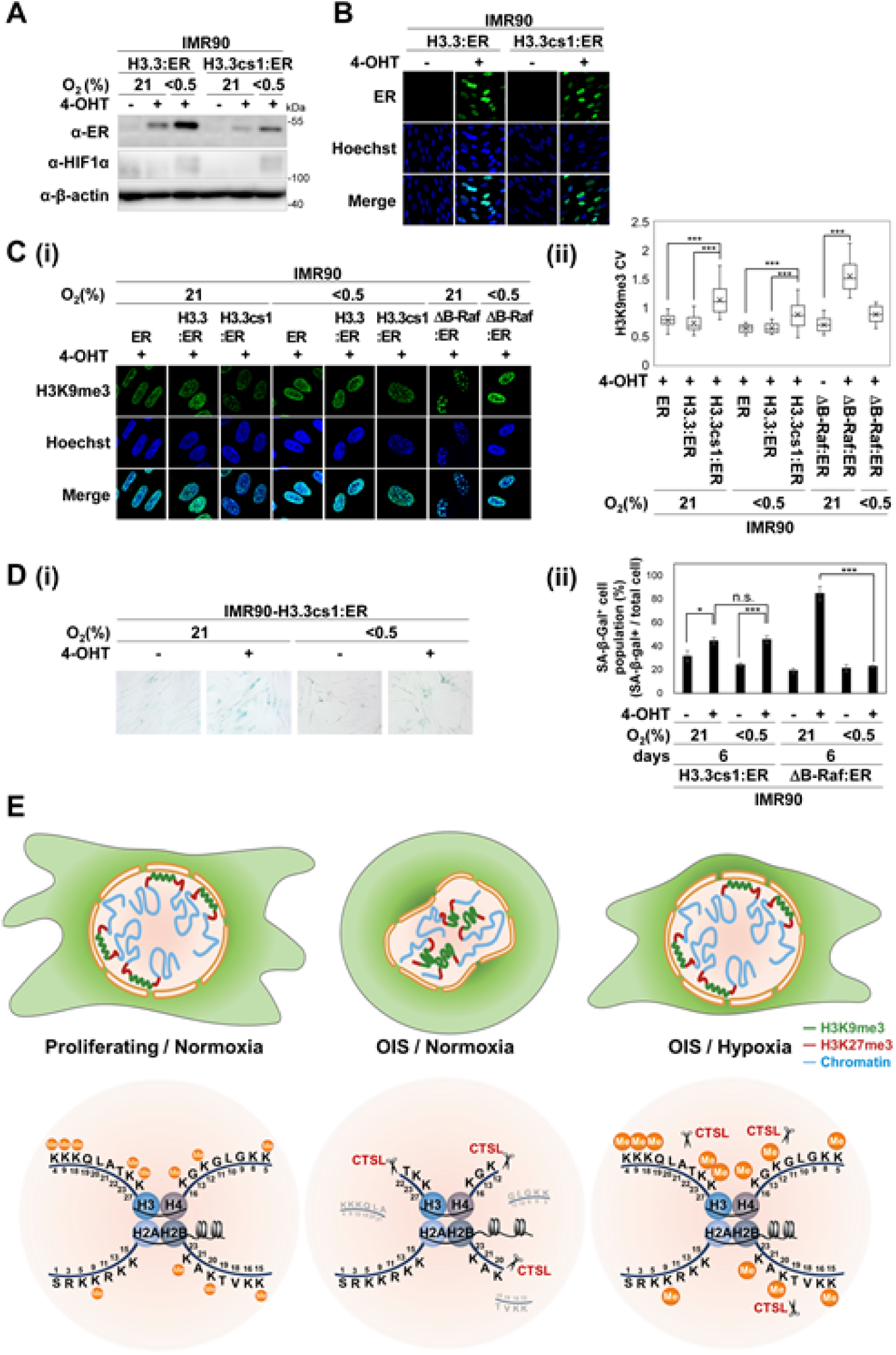
Forced expression of cleaved H3.3 (H3.3cs1) in hypoxic IMR90 cells. IMR90-ER, IMR90-H3.3:ER, and IMR90-H3.3cs1:ER cells are IMR90 cells infected with retroviruses encoding ER, H3.3:ER, and H3.3cs1:ER, respectively. The cells were treated as indicated for 6 days. (A) Western blot analysis. (B) Confocal images obtained following staining with ERα antibody and Hoechst. (C) (i) Confocal microscopic images of H3K9me3 and Hoechst staining of indicated cells. (ii) CV boxplots of images stained with the H3K9me3 antibody. H3K9me3 intensities in 20–35 nuclear images of each cell type were detected. (D) (i) Representative images of the SA-β-gal assays. (ii) Mean and S.D. of SA-β-gal-positive cells from four randomly selected microscopic fields. SA-β-gal-positive cells among 100–200 cells per field were counted. \**p* < 0.05, \*\**p* < 0.01, and \*\*\**p* < 0.001 by Students’ t-test; ns, not significant. (E) A schematic representation of hypoxic effects on histone clipping and SAHF during OIS (details in RESULTS).

## DISCUSSION

Hypoxic conditions in stem cell niches and inside solid tumors contribute to the inhibition of senescence and differentiation, thus maintaining the quiescent and undifferentiated status of resident cells. Hypoxia directly increases methylated histones by inhibiting the catalytic activities of O_2_-and α-KG-dependent histone lysine demethylases (JHDMs/KDMs). This study investigated how hypoxia-induced increase of methylated histones affects other epigenetic changes, such as histone clipping and redistribution of heterochromatin, which was found during terminal differentiation and OIS. The OIS process is a good model system for this study for the following reasons: Forced expression of oncogenes, such as Raf, in primary IMR90 cells triggers a variety of epigenetic changes, such as global histone demethylation, redistribution of heterochromatin, and especially histone clipping. Second, hypoxia has profound effects on the delay or prevention of OIS (58). Third, within a short period (4 to 6 days), various epigenetic changes can be observed. Fourth, the redistribution of heterochromatin during OIS, named SAHF, is so obvious that it can be observed in microscopic images stained with heterochromatin markers, such as H3K9me3, HP1, and H3K27me3.

By investigating the molecular mechanism by which hypoxia intervenes in epigenetic changes driven by OIS, we dissected pleiotropic and simultaneous events to deduce cause-and-effect relationships among events in nuclei during OIS. The decrease in methylated histones is necessary for OIS to trigger both histone clipping and SAHFs, but hypoxia maintains methylated histones to prevent both. Forced reduction of methylated histones in hypoxic cells using inhibitors of methyl-transferases makes hypoxic conditions unable to prevent OIS from inducing both histone clipping and SAHFs. Reversely forced enhancement of methylated histones using KDM pan inhibitors protected histones from clipping during OIS, even under normoxia, but failed to block SAHFs. Altogether, these results suggest that maintenance of methylated histones is sufficient to protect histones from CTSL, but is not sufficient but necessary for inhibiting SAHFs, suggesting that besides inhibiting KDMs, hypoxia adopts other methods to inhibit SAHFs.

Methylations of H3K4, H3K9, H3K27, and H3K36 have been found to decrease during OIS, which is caused by the induction of histone demethylases: KDM5/JARID1 (H3K4me1/2/3-demethylases) (59), KDM4/JMJD2 (H3K9me1/2/3 and H3K36me1/2/3-demethylases) (60), and KDM6/JMJD3 (H3K27me1/2/3-demethylases) (61,62). In addition, several methyl transferases, such as EZH2, a component of H3K27 specific methyltransferase complex, were found to be repressed during OIS (63). Here, we first found that both H3K23me3 and H3K18me3 levels were also reduced during OIS. The methylation of the H3K23 and K18 residues remains poorly understood. Methylated H3K23 was first identified in the heterochromatin region of Tetrahymena thermophila (53,64) and Caenorhabditis elegans (65). Recently, mass spectrometry revealed the presence of methylation and acetylation of H3K23 and H3K18 in human cancer cells (66). Point mutations in H3K23 and H3K18 residues have also been found in human cancers, but the contribution of these mutations to tumorigenesis remains unclear (54). Middle-down proteomic analyses (65) and ChIP-seq analyses (67) suggested that methylation of H3K23 positively correlates with heterochromatin histone marks, H3K9me3, H3K27me3, and heterochromatin proteins, but negatively correlates with H3K36me2/3 and H3K27ac, enriched in actively transcribed regions, suggesting that methylation of H3K23 contributes to organizing the genome in heterochromatin structure (68). In contrast, H3K23ac positively correlates with euchromatin markers (57). Enzyme kinetic analyses (Fig. 4D) and molecular dynamics simulations (Fig 4E) showed that tri-methylation of H3K23 directly reduced the fitness of H3 to CTSL, however it remains to investigate the indirect effects of H3K23me3 and H3K18me3 to CTSL. Histone peptide microarray analyses revealed that H3K23me2/3 specifically interact with chromodomains and double tudor domains, suggesting that H3K23me2/3 can recruit putative proteins which have specific chromo- or tudor-domains, to cover CTSL recognition sites (69,70). Compared to H3K23, the modification of H3K18 has barely been investigated. Methyltransferases and demethylases specific for both H3K23 and H3K18 residues are yet to be identified.

Using a polymer model of a chromosome, Sati et al. quantitatively assessed the mechanical strengths that determine SAHF formation during OIS (71). They found that two strengths, the interacting strength between nuclear lamina and heterochromatin and another interacting strength between heterochromatin domains should be reduced for SAHF formation. During OIS, the nuclear lamina has been found degraded (72,73). Our findings that hypoxia but not JIB-04 maintains the two strengths to prevent OIS from inducing SAHFs suggest that hypoxia may be related to other methods to inhibit SAHFs. It remains to investigate whether in addition to heterochromatin histone markers, hypoxia also maintains the integrity of nuclear lamina during OIS.

The cause-and-effect relationship between SAHF and chromatin accessibility remains unclear. Formation of SAHF results in redistribution of heterochromatin from the nuclear periphery to the inner large foci, causing decondensation of heterochromatin. This finding suggests that the OIS driven redistribution of heterochromatin (SAHF) may increase the accessibility of chromatin (71). The decondensed genomic regions encode senescence-up-regulated genes involved in cancer, organismal injury, abnormalities, and inflammatory responses (59,74). Cheung et al. first found that in human monocytes, three proteases, neutrophil serine proteases (NSPs), cathepsin G, and neutrophil elastase cleave N-terminus of H3 (13). They performed ChIP-seq analyses using an antibody specific for H3cs1 (cleaved H3 at T22) together with ATAC-seq and RNA-seq in human monocytes and revealed that the H3cs1 was enriched in open chromatin regions and at the TSS of permissive genes. Causality between histone clipping and chromatin accessibility was investigated. Simultaneous deletion of the three proteases in monocytes prevented H3 cleavage but increased chromatin accessibility at the TSS. These findings imply that proteolytic enzymes preferentially cleave histones in accessible open chromatin regions, and the cleaved histones then change chromatin architecture in a way that decreases chromatin accessibility. This idea is consistent with the finding that ectopic expression of H3.3cs1 decreases chromatin accessibility in human fibroblast IMR90 cells (22). Therefore, the increased accessibility of chromatin could be a cause, but not an effect, of histone clipping.

However, the role of cleaved histones in senescence remains unclear. This study consistently showed that ectopic expression of cleaved H3.3 (H3.3cs1) is sufficient to induce senescence (22). Differently from OIS, hypoxia could not prevent H3.3cs1 from inducing the senescence markers SA-β-gal and SAHFs, suggesting that histone clipping can reverberate senescence processes. We found that hypoxic conditions break off the positive feedback cycle by increasing histone methylation. These findings provide a novel mechanism by which hypoxia is involved in cell fate decisions by increasing methylated histones that protect histones and chromatin from dramatic epigenetic changes.

## DATA AVAILABILITY

Data that support the findings of this study are available from the corresponding author on reasonable request. Mass spectrometry data of histone top-down proteomics analyses are available at PRIDE database with the accession code PXD033971. Simulation protocols, input files and result files are available at zenodo.org as DOI: 10.5281/zenodo.6510402.

## Supporting information

manuscript and Figures

## ACKNOWLEDGEMENTS

We thank Prof. Martin McMahon (University of Utah) for providing plasmid of ΔB-RAF:ER. We thank Prof. Yun Doo Chung (University of Seoul) for providing Atto-590 Tn5.

