## Supplementary material for "Hypoxia increases the methylated histones to prevent histone clipping and redistribution of heterochromatin during Raf-induced senescence": manuscript and Figures

### Supplementary Figures

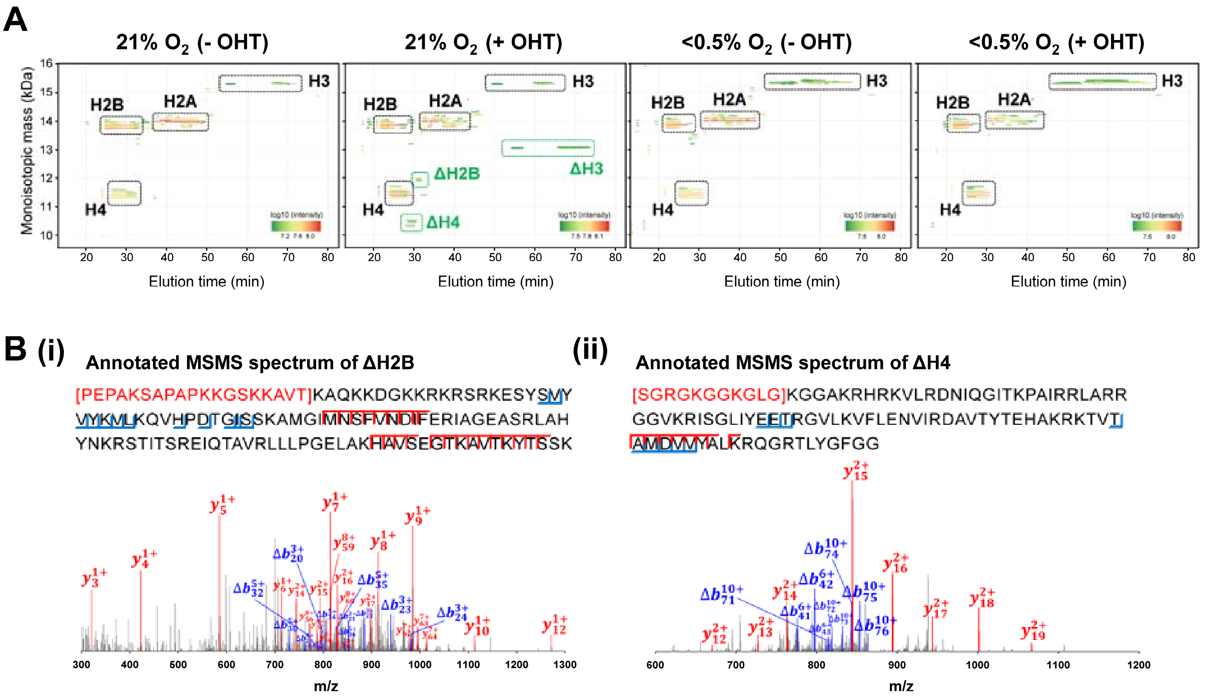


**Figure S1. Top-down proteomics of histone extracts.**

(A) Comparisons of 2D display of measured histones using samples obtained from IMR90-ΔB-RAF:ER cells treated with four different conditions (normoxia without OHT, normoxia with OHT, hypoxia without OHT, and hypoxia with OHT) between 15 and 80 min. (B) Annotated MS/MS spectra labeled with Δb ions (blue) and y ions (red) showing truncated H2B (i) and H4 (ii).

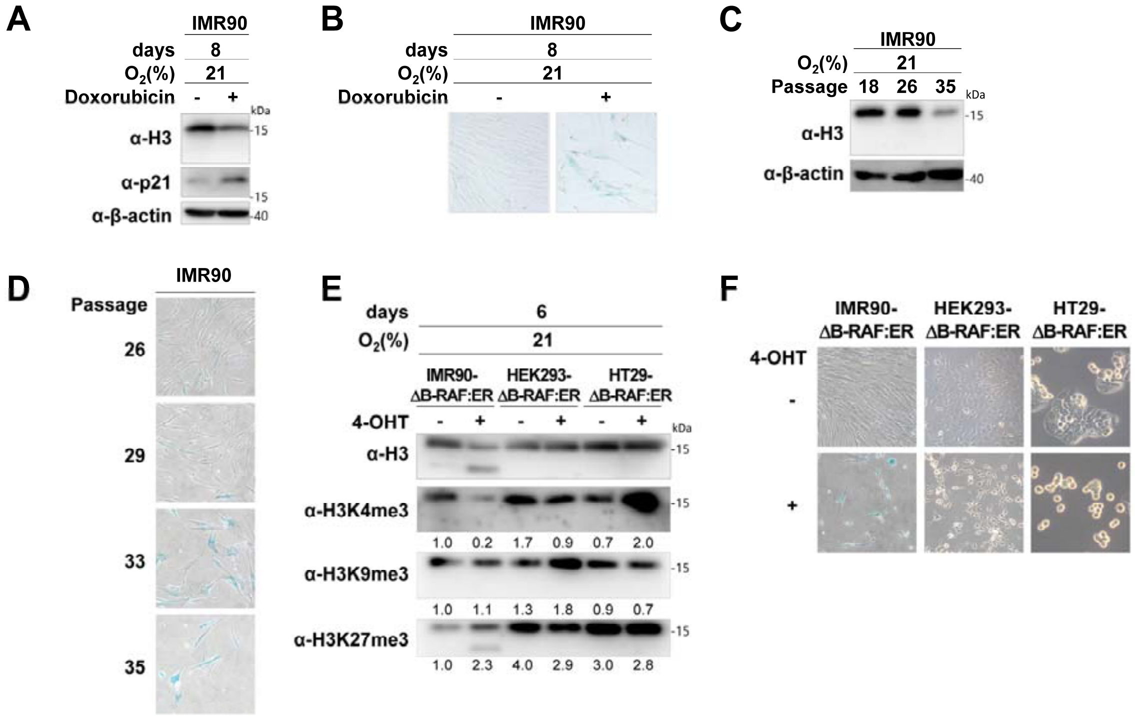


**Figure S2. Histone clipping during doxorubicin-induced senescence and replicative senescence.**

(A-B) Doxorubicin-induced senescence. IMR90 cells were treated with 100 nM doxorubicin for 2 days and then cultured in fresh media for an additional 6 days. (A) Western blot analysis using the indicated antibodies. (B) Representative images of SA-β-gal assay. (C-D) Replicative senescence induced by serial passaging of IMR90 cells as indicated. (C) Western blot analysis with the indicated antibodies. (D) Representative images of SA-β-gal assay. (E-F) IMR90, HEK293, and HT29 cells were infected with ΔB-RAF:ER and treated (+) with 4-OHT (100 nM) for 6 days. (E) Western blot analysis of histones using indicated antibodies. (F) Representative images of SA-β-gal assay.

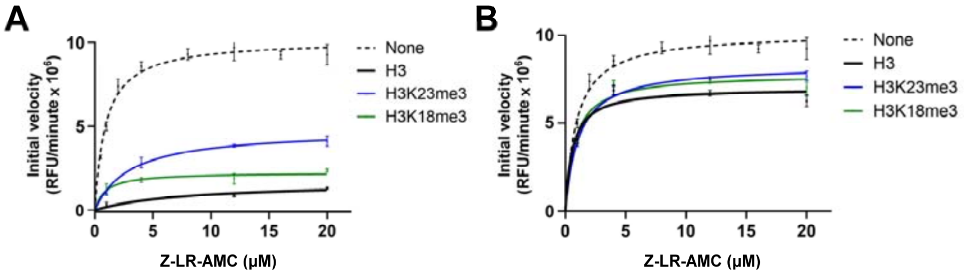


**Figure S3. Competitive inhibition assay of CTSL by H3 peptides with various modifications.** (A-B) Michaëlis-Menten plots of H3 peptides were fitted using an initial velocity with peptide concentrations of (A) 400 nM and (B) 100nM. The values on the y-axis indicate the mean and S.D. of initial velocity (RFU/min) of three replicates at given concentrations of Z-LR-AMC shown in x-axis. The Km and Ki values were determined using initial velocity and nonlinear regression fit with GraphPad Prism software from a representative single experiment among two independent experiments.

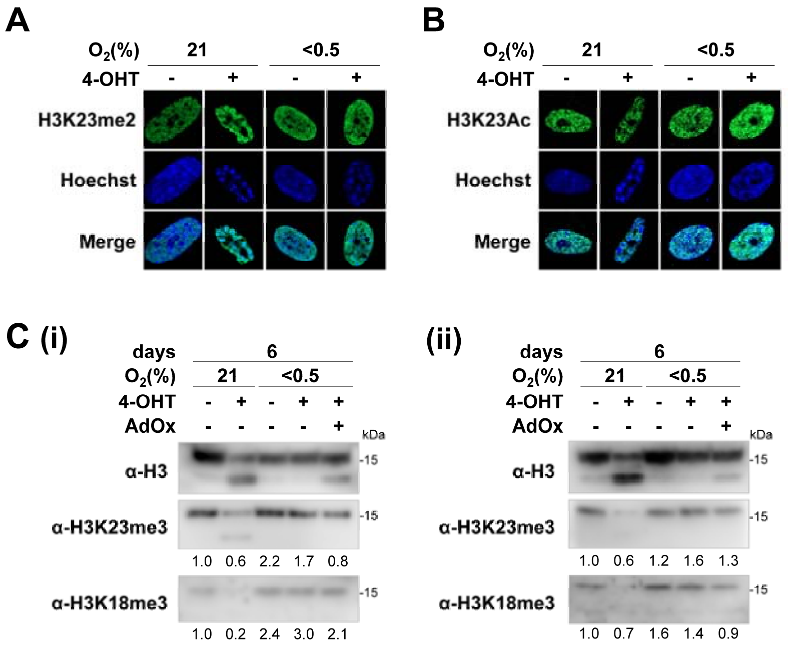


**Figure S4. H3K23me2 and H3K23Ac localization in OIS cells**

IMR90-ΔB-RAF:ER cells were treated for six days. Confocal microscopic images of nuclei stained with H3K23me2 (A), H3K23Ac (B) antibody, and Hoechst. (C) Western blot analysis with the indicated antibodies. The numbers represent the relative band intensities of methylated H3 normalized to the total H3 band intensities. Repeated results are shown in Fig. 5D.

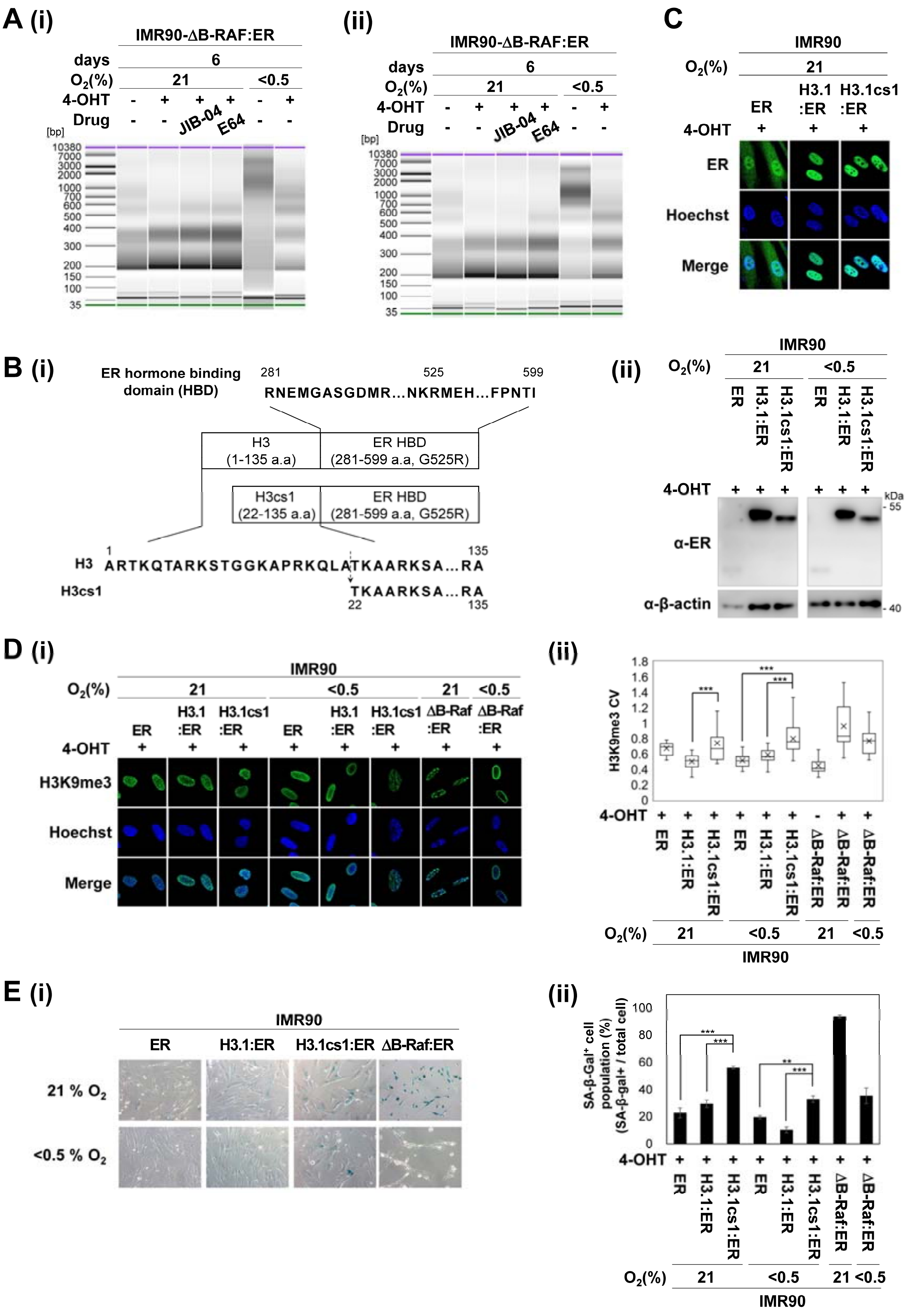

**Figure S5. Chromatin accessibility and forced expression of cleaved H3.1 (H3.1cs1) in hypoxic IMR90 cells.**

(A) Fragment sizes for amplified ATAC, as determined using a Bioanalyzer. The repeated results are shown in Fig. 6B. (B) (i) Schematic diagram of 4-OHT inducible H3:ER and H3cs1:ER fusion protein. (ii) Western blot analysis using the indicated antibodies. (C) Confocal images of the indicated cells stained with ERα antibody and Hoechst. (D) (i) Confocal microscopic images of cells stained with the H3K9me3 antibody and Hoechst. (ii) CV boxplots of images stained with H3K9me3 antibody. H3K9me3 intensities of total 20–30 nuclear images were counted. (E) (i) Representative images of the SA-β-gal assay. (ii) Mean and S.D. of SA-β-gal-positive cells from four to six randomly selected fields. SA-β-gal-positive cells among 50–85 cells per field were counted. *$p < 0.05$, **$p < 0.01$, and ***$p < 0.001$ by Students' t-test.

### Supplementary Tables

#### Table S1. Stable cells generated in this study

| Name | Host Cell | Gene | Vector / Selection marker |
|---|---|---|---|
| IMR90-ΔB-RAF:ER | IMR90 | ΔB-RAF:ER<br>B-RAF (449-804 aa) fused with hormone-binding domain of estrogen receptor (281-599 aa, G525R) | pBabe retroviral plasmid<br>/ Puromycin |
| HEK293-ΔB-RAF:ER | HEK293 | ΔB-RAF:ER<br>B-RAF (449-804 aa) fused with hormone-binding domain of estrogen receptor (281-599 aa, G525R) | pBabe retroviral plasmid<br>/ Puromycin |
| HT29-ΔB-RAF:ER | HT29 | ΔB-RAF:ER<br>B-RAF (449-804 aa) fused with hormone-binding domain of estrogen receptor (281-599 aa, G525R) | pBabe retroviral plasmid<br>/ Puromycin |
| IMR90-ER | IMR90 | ER<br>Hormone-binding domain of estrogen receptor (281-599 aa, G525R) | pBabe retroviral plasmid<br>/ Puromycin |
| IMR90-H3.1:ER | IMR90 | H3.1:ER<br>H3.1(1-135 aa) fused with hormone-binding domain of estrogen receptor (281-599 aa, G525R) | pBabe retroviral plasmid<br>/ Puromycin |
| IMR90-H3.1cs1:ER | IMR90 | H3.1cs1:ER<br>H3.1cs1(22-135 aa) fused with hormone-binding domain of estrogen receptor (281-599 aa, G525R) | pBabe retroviral plasmid<br>/ Puromycin |
| IMR90-H3.3:ER | IMR90 | H3.3:ER<br>H3.3(1-135 aa) fused with hormone-binding domain of estrogen receptor (281-599 aa, G525R) | pBabe retroviral plasmid<br>/ Puromycin |
| IMR90-H3.3cs1:ER | IMR90 | H3.3cs1:ER<br>H3.3cs1(22-135 aa) fused with hormone-binding domain of estrogen receptor (281-599 aa, G525R) | pBabe retroviral plasmid<br>/ Puromycin |
| IMR90-ΔB-RAF:ER-M56CTSL | IMR90-ΔB-RAF:ER | M56 CTSL-Myc-His<br>M56 CTSL (56-333 aa) fused with Myc-His tagged | pLenti CMV lentiviral plasmid<br>/ Hygromycin |
| IMR90-ΔB-RAF:ER EV | IMR90-ΔB-RAF:ER | None | pLenti CMV lentiviral plasmid<br>/ Hygromycin |
| IMR90-ΔB-RAF:ER-shCTSL | IMR90-ΔB-RAF:ER | shRNA against human CTSL<br>5’- GGTGGTTGGCTACGGATTTGA-3’ | pLKO.1 lentiviral plasmid<br>/ Hygromycin |
| IMR90-ΔB-RAF:ER-shScramble | IMR90-ΔB-RAF:ER | Scrambled shRNA<br>5’- CCTAAGGTTAAGTCGCCCTCG-3’ | pLKO.1 lentiviral plasmid<br>/ Hygromycin |

**Table S2. Antibodies.**

| Antibody against | Provider | Cat.No. | Application |
|---|---|---|---|
| ERα | Santa Cruz Biotechnology | sc-543 | WB, IF |
| p21 | Santa Cruz Biotechnology | sc-6246 | WB |
| HIF-1α | BD bioscience | 610959 | WB |
| 14-3-3γ | Millipore | 05-639 | WB |
| β-actin | Sigma | 3700 | WB |
| Cathepsin L | Abcam | ab6314 | WB, IF |
| Myc | IG therapy | A200302 | IF |
| His | Bethyl | A190-114A | WB |
| H2A | Abcam | ab18255 | WB |
| H2B | Abcam | ab1790 | WB |
| H4 | Abcam | ab10158 | WB |
| H3 | Abcam | ab1791 | WB |
| H3.1/3.2 | Millipore | ABE154 | WB |
| H3.3 | Millipore | 09-838 | WB |
| H3cs1 | Active motif | 39573 | WB |
| H3K4me3 | Cell signaling | 9751S | WB, IF |
| H3K9me3 | Abcam | ab8898 | WB, IF |
| H3K27me3 | Millipore | 07-449 | WB, IF |
| H3K79me2 | Cell signaling | 9757 | WB |
| H3K23me3 | PTM bio | PTM648 | WB, IF |
| H3K23me2 | Abcam | ab214654 | WB |
| H3K23Ac | Abcam | ab61234 | WB |
| H3K18me3 | Novus bio | NB21-1143 | WB, IF |
| H3K18me2 | Novus bio | NB21-1142 | WB |
| H3K18Ac | Abcam | ab40888 | WB |
| HP1β | Cell signaling | 8676P | WB, IF |
| Alexa Fluor ® 488 | Invitrogen | A11034 | IF |
| Alexa Fluor ® 546 | Invitrogen | A11030 | IF |

**Table S3. Reagents.**

| Reagents | Provider | Cat.No. | Final concentration, /Stock concentration |
|---|---|---|---|
| Hoechst 33258 | Sigma | 94403 | 1 μg/ml / 1mg/ml in PBS-T |
| JIB-04 | Sigma | SML0808 | 0.5 μM / 1mM in DMSO |
| E64 | Sigma | E3132 | 10 μM / 10mM DMSO |
| Adenosine dialdehyde (AdOx) | Sigma | A7154 | 50 μM / 50mM in DMSO |
| Ciclopirox (CPX) | Sigma | SML2011 | 10 μM / 10mM in Water |
| Deferiprone (DFP) | Sigma | 379409 | 250 μM / 10mM in Water |
| Dimethyloxalylglycine (DMOG) | Sigma | D3695 | 0.5 mM / 100mM in DMSO |
| Doxorubicin | Sigma | D1515 | 100nM / 100μM in Water |
| 4-hydroxy-tamoxifen (4-OHT) | Sigma | H7904 | 100nM / 1mM in EtOH |
| 3-deazaneplanocin A (DZNep) | Selleckchem | S7120 | 10 μM / 10mM in DMSO |
| Magic Red Cathepsin L Assay Kit | ImmunoChemistry Technologies | 941 | 3.6 μM / in DMSO [4] |
| Recombinant Human Cathepsin L | R&D systems | 952-CY-010 | 50pg/rxn[1] /1ng/μl in reaction buffer[3]<br>96pM[2] / 2μM in Reaction buffer[3] |
| Recombinant H3 | Active motif | 31294 | 200ng/rxn |
| Recombinant H2B | Active motif | 31492 | 200ng/rxn |
| Z-LR-AMC | R&D systems | ES008 | 1-20μM /32mM in DMSO |
| $H3_{14\text{-}31}$ Peptide (N-terminal biotinylated) | GL Biochem | Customized Peptide | 100nM, 400nM / 1mg/ml in Water |
| $H3_{14\text{-}31}$ K23me3 Peptide (N-terminal biotinylated) | GL Biochem | Customized Peptide | 100nM, 400nM / 1mg/ml in Water |
| $H3_{14\text{-}31}$ K23Ac Peptide (N-terminal biotinylated) | GL Biochem | Customized Peptide | 100nM, 400nM / 1mg/ml in Water |
| $H3_{14\text{-}31}$ Peptide | EZBiolab | Customized Peptide | 100nM, 400nM / 1mg/ml in Water |
| $H3_{14\text{-}31}$ K23me3 Peptide | EZBiolab | Customized Peptide | 100nM, 400nM / 1mg/ml in Water |
| $H3_{14\text{-}31}$ K18me3 Peptide | EZBiolab | Customized Peptide | 100nM, 400nM / 1mg/ml in Water |
| Tn5-Nextera DNA library prep kit | Illumina | FC-121-1030 | TD buffer: 25 μL/rxn [4]<br>TDE1 enzyme: 2.5 μL/rxn [4] |
| Nextera Index Kit | Illumina | FC-121-1011 | 5 μL/rxn [4] |
| Next High-Fidelity 2× PCR Master Mix | NEB | M0541S | 25 μL/rxn [4] |
| Atto-590 Tn5 | Dr.Yun Doo Chung (University of Seoul) | N/A | 1 μL/rxn |

[1] *In vitro* proteolytic reaction

[2] Kinetic assay

[3] Reaction buffer (50 mM MES (pH 5.5), 5 mM DTT, 1 mM EDTA, and 0.005% Brij-35.)

[4] Reagents provided as a solution. We used vol/rxn as guided.
